# Haplomatic: A Deep-Learning Tool for Adaptively Scaling Resolution in Genetic Mapping Studies

**DOI:** 10.1101/2025.06.25.661582

**Authors:** Tyler Douglas, Anthony Long, Rebecca Tarvin

## Abstract

Genomic mapping studies face a fundamental trade-off between accuracy and resolution: increasing resolution improves localization of genetic signals but typically reduces the accuracy of frequency estimates due to increased statistical noise. In pooled sequencing studies, this trade-off impacts the accuracy of haplotype frequency estimates, a primary statistic used to identify genetic associations.To mitigate this trade-off we introduce Haplomatic, a novel deep-learning-based tool that adaptively adjusts genomic resolution by predicting haplotype frequency estimation error. Haplomatic generates simulated population data from known recombinant inbred line populations, predicts error through a transformer-based neural network, and adjusts resolution until a target error is achieved. Haplomatic achieves significant resolution gains (15% on average) over previous methods without sacrificing accuracy across multiple evaluated sequencing depths (10x, 50x, and 100x). To our knowledge, this is the first instance of applying deep learning to directly predict estimation error and dynamically scale resolution in trait mapping studies.

## Introduction

A fundamental trade-off between resolution and statistical reliability exists in genomic mapping studies: finer resolution facilitates precise localization of genetic signals but increases noise, obscuring actual associations and potentially generating false positives (Mackay et al., 2009; Kofler and Schlötterer, 2014). This trade-off is especially pronounced in studies using pooled sequencing, where quantitative trait loci (QTL) are identified by associating phenotypic variation with SNP frequency estimates at the population level (Long et al., 2015; Schlötterer et al., 2015). In these systems, noise in SNP frequency estimates limits mapping accuracy and can complicate candidate gene identification. Aggregating SNP data into genomic windows and inferring haplotype frequencies within these windows rather than relying on individual SNP frequencies can mitigate the effects of noise in SNP frequency estimates (MacDonald et al. 2022). However, selecting an optimal window resolution is challenging: smaller windows increase resolution but risk higher estimation errors due to insufficient informative SNPs, whereas larger windows enhance estimation accuracy but obscure fine-scale genomic features (Franssen et al., 2017; Barghi et al., 2020).

Previous methodologies have typically employed fixed, genome-wide window sizes based on physical distance (200 kb and larger) or genetic distance (1.0 cM), although this does not account for local variation in SNP density and informativeness (Turner et al., 2011; Jha et al., 2015).

Deep learning is increasingly being applied to genetic mapping studies to improve identification of causal loci. Tools such as DeepVariant, which uses convolutional neural networks to improve variant calling, or SuSie, which uses Bayesian methods to identify causal variants focus on SNP-level inference (Poplin et al. 2018; Wang et al. 2020). However, no current method dynamically adapts mapping resolution based on predicted estimation accuracy or directly quantifies this error to optimize resolution genome-wide. And, none of the above tools are compatible with pooled sequencing data.

To address this gap we developed Haplomatic, a deep-learning-based tool that adaptively scales genomic resolution in pooled sequencing trait mapping studies. Haplomatic simulates in silico populations derived from known recombinant inbred line (RIL) panels such as the Drosophila Synthetic Population Resource (DSPR; King et al., 2012), uses a transformer-based neural network to predict haplotype frequency estimation error, and dynamically adjusts resolution to achieve a specified accuracy target. This approach leverages deep learning to directly predict estimation errors, enabling adaptive genomic resolution based on local genomic characteristics. To our knowledge, Haplomatic represents the first attempt to apply deep learning techniques for direct error prediction and dynamic resolution adjustment in trait mapping. By enhancing resolution without compromising accuracy, Haplomatic can significantly improve the precision of genomic mapping from evolve-and-resequence experiments.

## Methods

### Population Simulation

To generate training data, we simulated 20 *in silico* populations consisting of 300 individuals by sampling random recombinant inbred lines (RILs) with replacement from an initial pool of 500 Drosophila Synthetic Population Resource (DSPR) RILs. DSPR founder genotypes of the starting RILs were resolved at 10kb intervals by individual sequencing. After randomly selecting 300 lines, each initial population underwent forward simulation for 10 subsequent generations, with individuals resampled with replacement each generation to simulate drift. Recombination events were introduced by randomly exchanging chromosome segments at 10 kb intervals, at a rate of 0.5 events per chromosome per generation. This produced simulated populations with known haplotype frequencies, which we used both for model training and to calculate correlations between accuracy and various local genomic features (SNP number etc.).

### Read Simulation

To simulate paired-end reads from the evolved populations, we first constructed DSPR founder fasta sequences of chromosome 3L by projecting homozygous SNPs from each DSPR founder line onto the *D. melanogaster* Release 6 reference genome using GATK’s FastaAlternateReferenceMaker (Hoskins et al. 2015; McKenna et al. 2010). Sequencing data was then simulated using Haplomatic at 100x coverage from each of the five evolved populations by randomly selecting genomic start positions along chromosome 3L and sampling read lengths from a normal distribution (mean = 300 bp, standard deviation = 50 bp). Actual read sequences were derived from the corresponding founder reference fasta at the selected read positions.

Simulated reads were mapped to the *Drosophila melanogaster* Release 6 using bwa-mem2 (v2.2.1; Li 2013), and 100x data was then downsampled to 50x and 10x coverage. Variant calling was performed using bcftools (v1.17; Danacek et al. 2021). Finally, a SNP frequency table was generated containing the frequency of alternate allele in the 8 DSPR founders and simulated populations

### Haplotype Frequency Estimation

Within each genomic window, the observed SNP frequency vector *b* was modeled as a linear combination of founder haplotypes with Gaussian noise. Given *X*, a binary matrix indicating the presence or absence of alternate alleles in each founder haplotype across the SNPs in the window, we defined the model as:

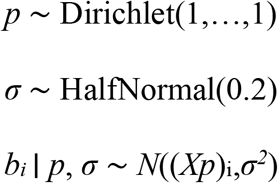

where *p* represents the haplotype frequency vector to be inferred, and *σ* represents observation noise. Starting from a Dirichlet prior ensured haplotype frequencies must sum to one. Inference was performed using the No-U-Turn Sampler (NUTS) in NumPyro (Phan et al. 2019). For each window, we ran one MCMC chain with 100 warm-up iterations and 100 sampling iterations.

Posterior samples of *p* were summarized by their means and were also used to compute model features including posterior standard deviation and skewness.

### Model Features and Architecture

We calculated model features by splitting the SNP frequency table described above by SNP position into windows of various lengths spanning 30 kb to 250 kb. Within each window, we calculated a feature set using the simulated variant call data to summarize local genomic characteristics (Table 1). Each window was then assigned a corresponding true haplotype frequency vector, identified by locating the midpoint of the window and retrieving the known true frequencies from the corresponding simulated population. Finally, we calculated estimation error for each window as the total absolute difference between true and inferred haplotype frequencies and defined this as our training target.

**Table 1.**
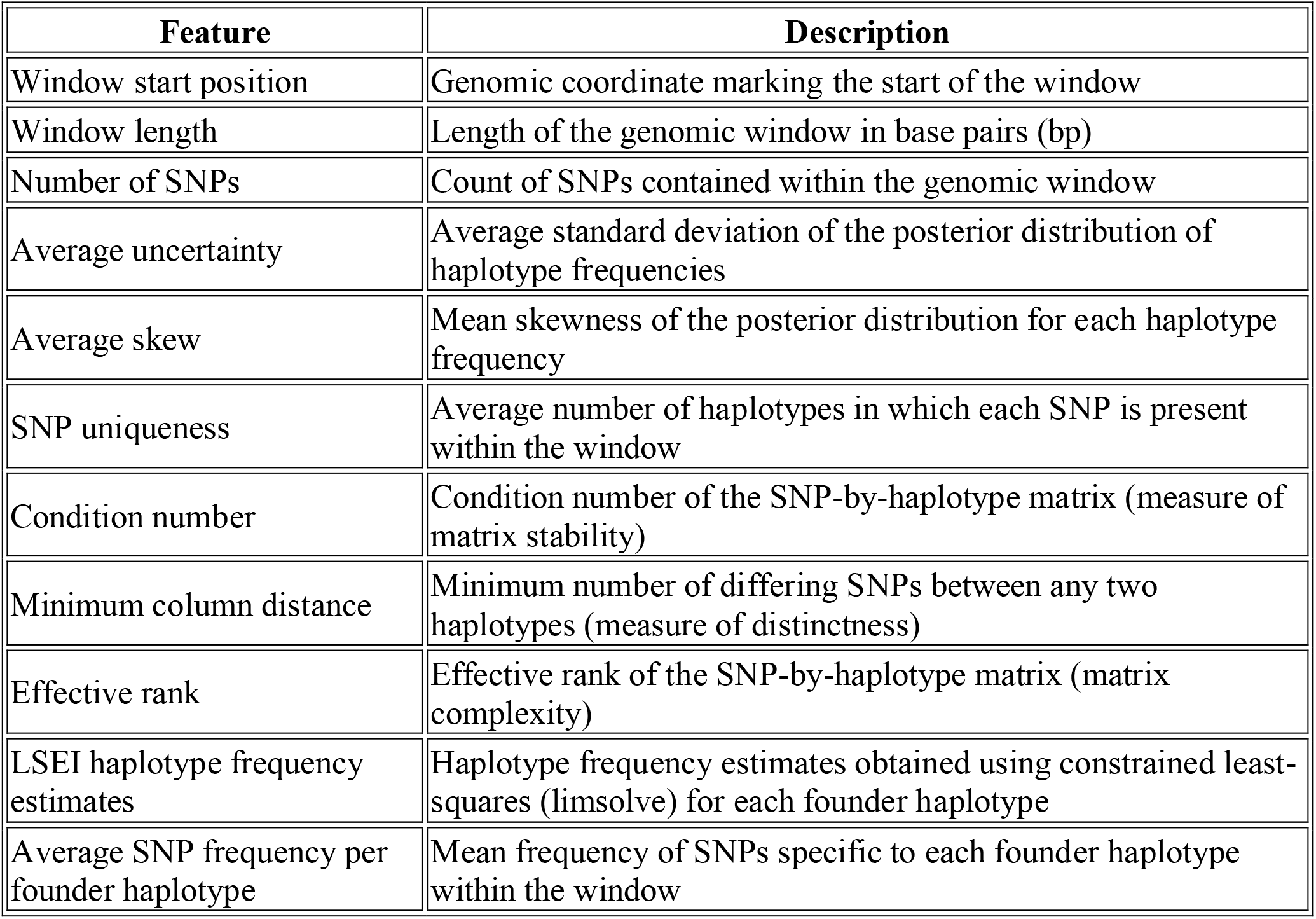
Model features calculated per genomic window.

To predict estimation error, we utilized a hybrid neural network in Pytorch v2.7 consisting of two main components: a transformer-encoder that processes all raw SNPs in a genomic window from the SNP frequency table, and a feedforward regressor that processes our window summary features (Table 1; Paszke et al. 2019). Specifically, the transformer-encoder consisted of a linear embedding layer (embedding dimension = 64), followed by three stacked transformer-encoder layers with four attention heads each. A learned attention-weighted pooling layer aggregated encoded SNP-level representations into a fixed-length vector. This output was concatenated with the feature vector of local genomic characteristics (Supplementary Table 1) and passed to a regressor consisting of two hidden layers (256 and 128 units) with ReLU activations and dropout (rate = 0.2) applied after each layer.

### Model Training

Each training instance corresponded to a genomic window and included three components: a SNP matrix *X* of shape *n* SNPs × *k* founders encoding the presence (0/1) of each SNP in each founder; a vector of window summary features, and a scalar target representing the total absolute error between true and inferred haplotype frequencies for that window. To standardize input size for the transformer encoder, SNP matrices were zero-padded or right-cropped to 400 SNPs per window.

Training was performed in three phases. A model was first trained for 40 epochs using data from 15 simulated populations at 10x sequencing coverage. This model was then fine-tuned for an additional 20 epochs using simulated populations at 50x and 100x coverage. Subsequently, for each coverage level, specialist models were trained on medium and low error windows specifically, to capture more fine-grained patterns than those captured in a single global model. The remaining five populations at each coverage level were held out and used exclusively for model validation. For all training runs, data were split 70/30 into training and validation sets, and tabular features were standardized using a standard scaler fitted on the training set. The model was optimized using Adam (learning rate = 1e-3, weight decay = 1e-5) with mean squared error loss. Training was performed with a batch size of 64, and the model with the highest validation R^2^ was saved.

### QTL Scans on Real Populations

Generation and artificial selection of experimental populations is described in Hanson et al. 2025. To identify genomic loci significantly associated with zinc resistance, we modeled arcsine-transformed haplotype frequencies as a function of treatment x haplotype with replicate as a random effect. We fit an ANOVA at each position and identified regions where the log-transformed significance of treatment x haplotype exceeded 4.7 (threshold set based on Bonferroni correction). To account for noise between windows, we LOESS-smoothed -log_10_(P) values (span = .04).

## Results and Discussion

### Accuracy of Bayesian Frequency Estimates

We characterized the accuracy of Bayesian haplotype frequency estimates across 10–250 kb windows at 10x, 50x, and 100x coverage by measuring the mean error per haplotype frequency estimate across the eight DSPR founder haplotypes. Mean error declined with increasing window size and coverage, achieving 3.7%, 3.1%, and 3.0% accuracy at 60 kb windows for 10x, 50x, and 100x data respectively. While average error continued to decrease as window size exceeded 60kb, we observed diminishing returns for comparatively large costs in resolution (Fig. 1A) and a weak relationship between window size and error (R^2^ = 5.5%). The observed distributions of error (Fig. 1B) indicate that while an average resolution of 60 kb yields acceptable accuracy overall, prediction error remains variable across regions: some regions can support higher resolution, whereas others require larger windows to maintain accuracy. In some genomic regions, increasing the window size actually led to higher estimation error, likely due to the inclusion of SNPs that blur distinctions between similar haplotypes and introduce conflicting signals. Two other intuitive predictors of error, namely SNP density and minimum column distance–a measure of how SNPs distinguish haplotypes–were also weakly related to estimation error (R^2^ = 7.8% and 7.7% respectively). These results suggest that estimation error is a complex, non-linear interaction between multiple weak predictors, including haplotype composition and local SNP informativeness.

**Figure 1.**
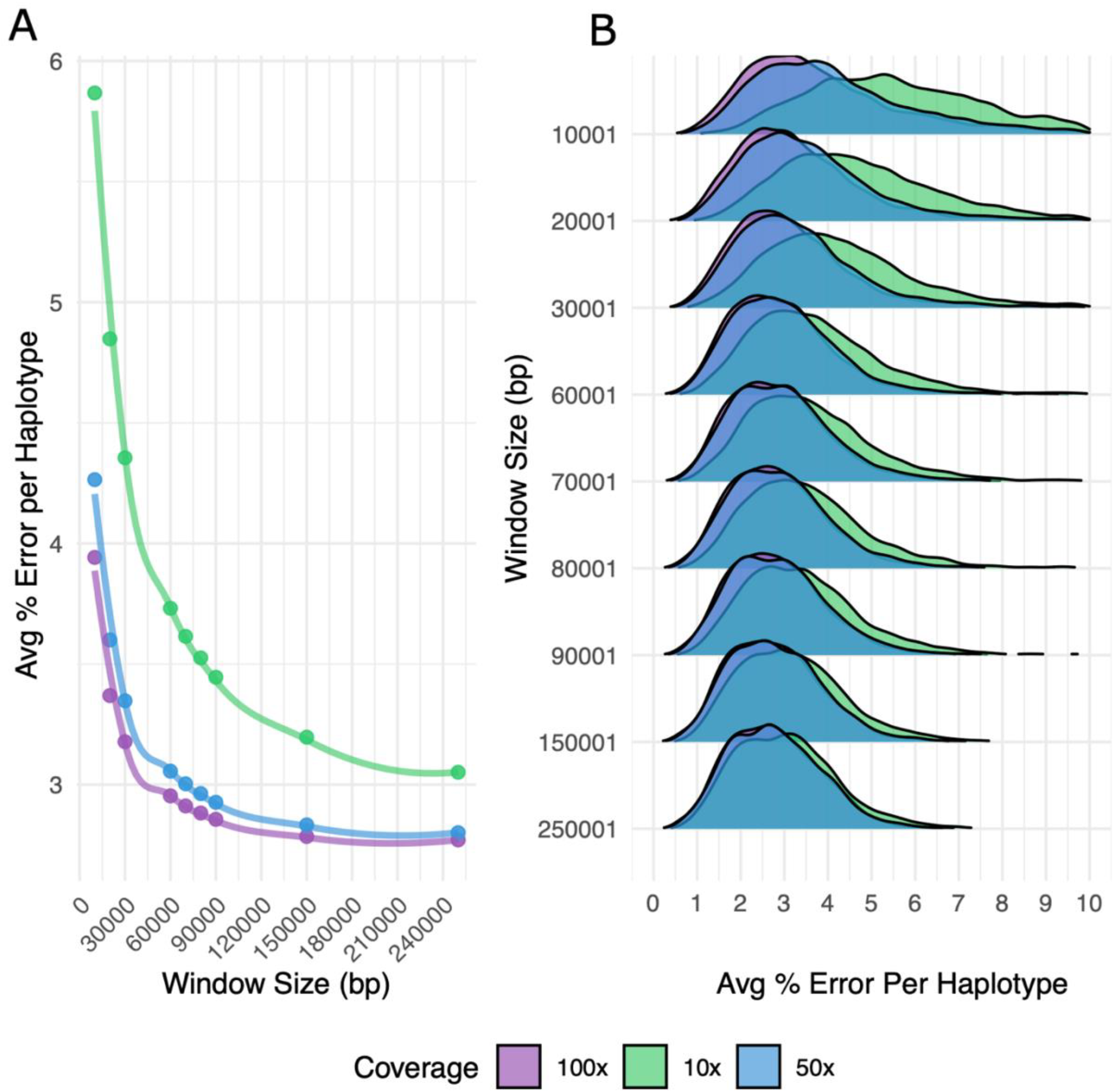
Accuracy of Bayesian haplotype frequency estimates across window sizes. A) Mean per-haplotype error as a function of window size, colored by sequencing coverage (10x, 50x, 100x). B) Distribution of per-haplotype error values at each window size colored by coverage.

Moreover, while it is often assumed that increasing sequencing coverage uniformly improves accuracy, we found that this advantage is limited to smaller window sizes. However, uniformly applying a small window size risks elevated estimation error even in high coverage datasets (Fig. 1B). While increasing window size can mitigate this risk, doing so diminishes the returns of sequencing at high coverage. For instance, at a window size of 250 kb, haplotype frequency estimates from 100x data were only 0.28% more accurate on average than those from 10x data. Therefore, adaptively scaling window size is necessary to realize the benefits of deeper sequencing without inflating error.

### Benchmarking Against Fixed Resolution

Previously published methods of haplotype frequency estimation have utilized a fixed physical distance of 200 kb or genetic distance of 1 cM to estimate genome-wide haplotype frequencies using a least-squares approach in *limsolve* (Soetaert et al. 2009). However, we observe that high accuracy estimates are sometimes possible at much lower resolutions (Fig. 1B). We assessed the gains in resolution and costs in error of haplotype frequency estimates from an adaptive resolution relative to fixed windowing. Specifically, we compared the accuracy (Fig. 3A) and window size (Fig. 3B) of an adaptive scan set to a target error of 2.5% to data from a fixed resolution of 200 kb, and a high resolution of 50 kb. As not all genomic regions conform to the minimal error threshold, we applied a “choose best” approach when predicted error failed to reach target error.

At 100x coverage, our adaptive model achieved essentially the same mean accuracy as a conservative fixed resolution (2.3% error at fixed 200 kb vs. 2.4% with adaptive resolution) but increased resolution by an average of 56.6% compared to the fixed 200 kb window. Using a fixed high resolution of 50 kb slightly raised the mean error (to 2.6%) but, more importantly, produced spurious error peaks exceeding 5% per haplotype error (40% total error) throughout the genome and particularly in centromeric regions, which can introduce false signals in downstream mapping. In contrast, the adaptive approach eliminated the artifacts generated by a fixed 50 kb resolution, and error only rose in regions where high error was unavoidable regardless of window size, as similar spikes in error occurred in those regions at the 200 kb scale. Notably, the adaptive model achieved resolutions finer than 50 kb across 13% of the chromosome, providing substantial stretches of high resolution where the data supported it. At lower coverages (50x and 10x), resolution gains remained meaningful (43% and 44% respectively) with no appreciable loss in accuracy. This was true even on low-coverage data, demonstrating that Haplomatic can maximize resolution even when sequencing costs must be minimized.

### Accuracy of Error Prediction

Adaptively scaling resolution by region depends on reliable predictions of accuracy. We found that models trained on 10x, 50x, and 100x coverage data predicted the error of haplotype frequency estimations on holdout datasets with an R^2^ of 72%, 75%, and 77% respectively. To directly assess Haplomatic’s ability to achieve a specified target accuracy, we performed haplotype frequency scans of the *D. melanogaster* chromosome 3L in 5 simulated hold out populations. For each scan, we specified a per-haplotype target accuracy of 2%-5%, (equivalent to 16–40% total error across all eight haplotypes). Some genomic regions exhibit high error irrespective of window size and do not conform to low error thresholds. Because our aim was to assess each model’s ability to identify windows of a specific error, in this scan we strictly included only genomic regions where predicted error was less than the specified target error. We found that, for 50x and 100x coverage, average error reliably falls within 0.5% of the specified target error across our range of tested target errors (Fig. 2A). Although this pattern held for 10x data, for target errors greater than 3%, a lower target error was not feasible, with 3% being the lowest achievable mean error on low coverage data. Further, both 50x and 100x data yielded an average per haplotype error of 2.3% with an average resolution of approximately 62 kb, while 10x data required a resolution of 110 kb to produce estimates within 3% accuracy. Thus, Haplomatic yields frequency estimates that match a user-defined target accuracy, except when low target error is infeasible given sequencing depth.

**Figure 2.**
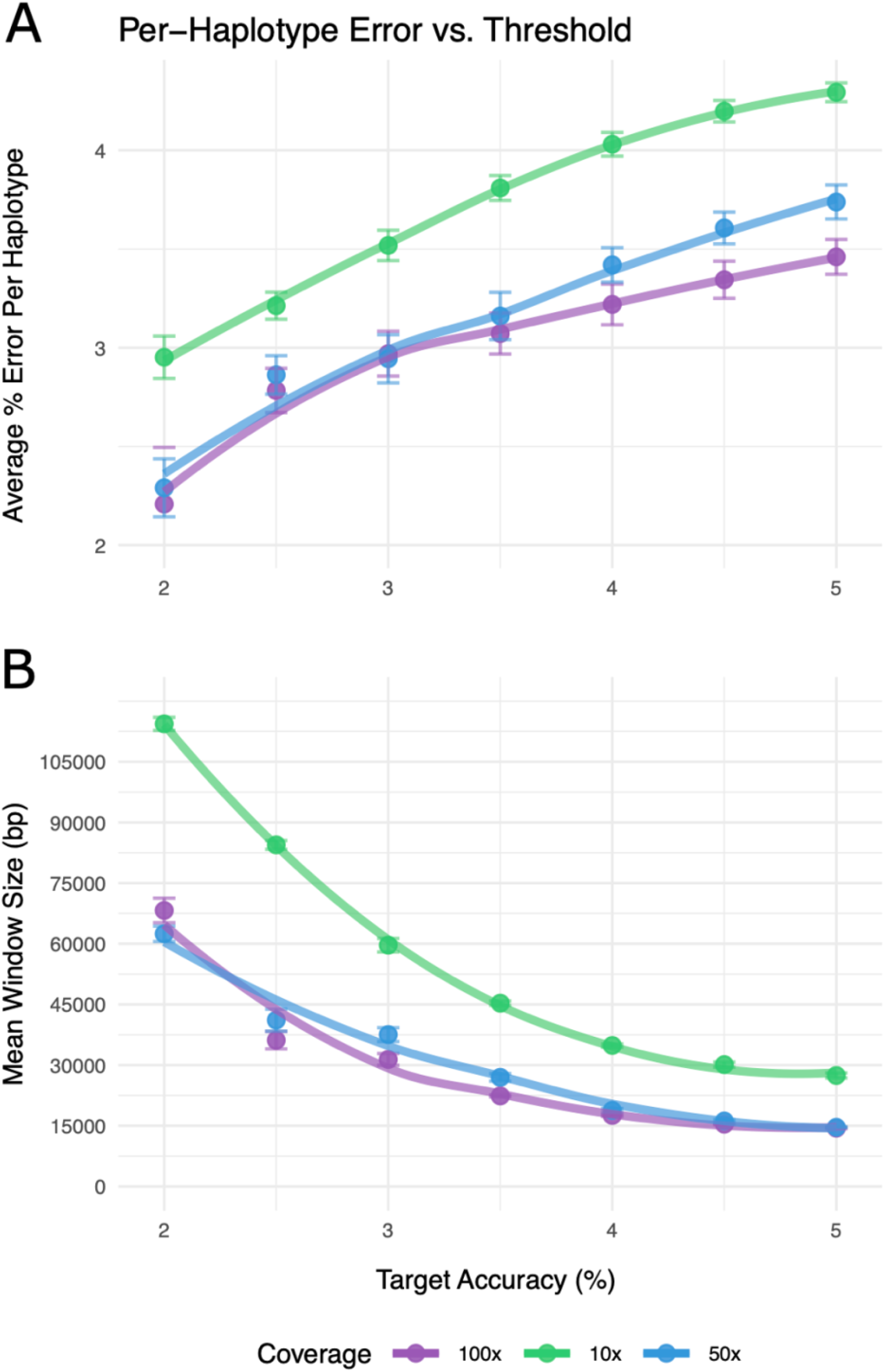
Model performance in achieving target estimation error. A) Actual per-haplotype error versus specified target error across simulations. B) Average genomic resolution (window size in bp) used to achieve each target error. Lines are colored by sequencing coverage level.

**Figure 3.**
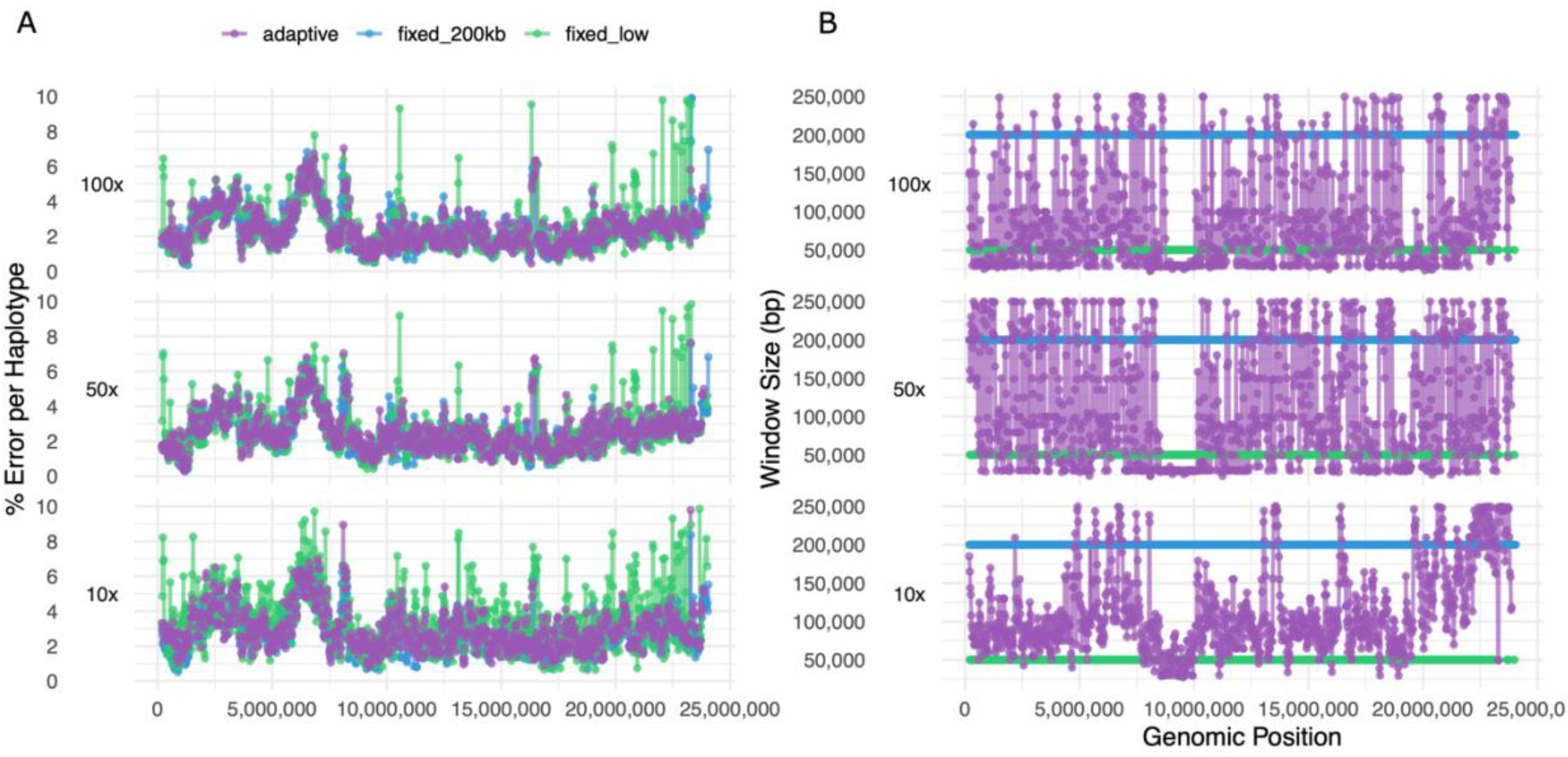
Genome-wide haplotype frequency scan along *Drosophila melanogaster* chromosome 3L. A) Estimated haplotype frequency error across the chromosome using adaptive windowing (target error = 2.5%) and fixed 50 kb and 200 kb windows. B) Corresponding window sizes used by each method. Rows correspond to 100x, 50x, and 10x sequencing coverage.

Notably, Haplomatic uses basic genomic features (for example SNP number and density, allele frequency distributions) and posterior distribution shape to predict estimation error. Our model does not depend on species-specific genomic annotation or species-specific feature engineering. Therefore, while we tested Haplomatic on data from *Drosophila melanogaster*, we expect Haplomatic to generalize well to other model systems.

### Validation on Real Populations

Haplotype frequency estimations over narrower genomic intervals should yield better QTL localization. To assess improvement in QTL resolution, we applied Haplomatic to actual data from DSPR-established populations of *D. melanogaster* that underwent selection for zinc resistance (Hanson et al. 2025). Specifically, we estimated haplotype frequencies across chromosome 3L at fixed (50 kb and 200 kb) and adaptive resolutions, and identified peaks where -log_10_(P) exceeded 4.5. We identified 4 prominent QTLs at 200 kb and 5 in the adaptive scan (Fig. 4), as Haplomatic split one of the QTLs in the fixed scan into two separate peaks.

**Figure 4.**
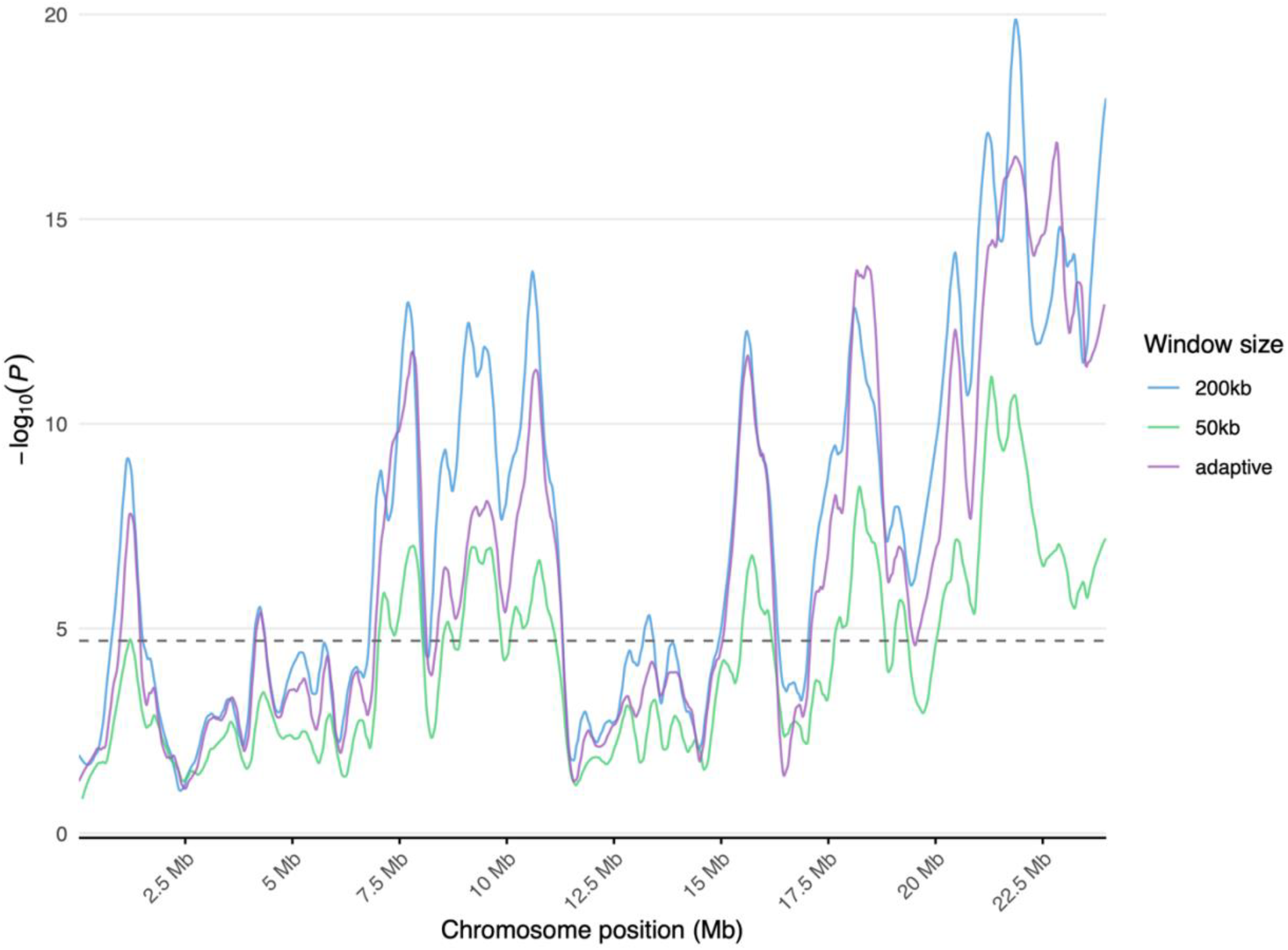
QTL scan of chromosome 3L from zinc-adapted *Drosophila melanogaster* populations.

Additionally, the adaptive scan yielded a chromosome-wide increase in haplotype frequency estimate resolution of 30% (140 kb versus 200 kb), and an average decrease in QTL peak width of 14.5%. In contrast, our 50 kb scan of chromosome 3L lacked sufficient power to detect QTL 1 and at times broader, flatter peaks, potentially due to increased noise in haplotype frequency estimates taken at an excessively high resolution. Taken together with our benchmarks on simulated populations, these results suggest that Haplomatic can yield an average increase in QTL resolution by an order of magnitude without costs to accuracy.

### Limitations

While Haplomatic can yield substantial gains in resolution, we note that it is much more computationally intensive than previous methods based on *limsolve*. Haplotype frequencies can be estimated along an entire chromosome in a few minutes on a local machine using *limsolve*, yet the same region takes approximately 4-5 hours using Haplomatic. Given its computational costs, Haplomatic is most well-suited not as a complete replacement for *limsolve*-based approaches, but as a follow-up to achieve higher resolution in targeted areas identified by an initial pass.

## Conclusion

Haplomatic provides a flexible solution to the trade-off between accuracy and resolution in pooled sequencing-based trait mapping. By leveraging deep learning to predict estimation error, it adaptively adjusts window sizes to maintain target accuracy thresholds, outperforming fixed-window approaches in both resolution and error control. Our results demonstrate that Haplomatic can substantially improve haplotype frequency resolution—achieving up to 13% higher resolution on real data without increasing error—while avoiding spurious signals introduced by overly narrow fixed windows. This framework offers a practical solution for evolve-and-resequence experiments and other genomic mapping efforts where pooled sequencing data and known founder haplotypes are available.

## Code and Data Availability

Haplomatic is available as an open-source Python package at https://github.com/tyleredouglas/haplomatic.

